# Seasonal microbial dynamics on grapevine leaves under biocontrol and copper fungicide treatments

**DOI:** 10.1101/523977

**Authors:** Gobbi Alex, Kyrkou Ifigeneia, Filippi Elisa, Ellegaard-Jensen Lea, Hansen Lars Hestbjerg

**Affiliations:** Environmental Microbial Genomics (EMG), Aarhus University, Department of Environmental Science, Roskilde, DK, Denmark

**Keywords:** Biocontrol, Fungicide, Grapevine, Meta-barcoding, qPCR, Copper, Amplicons, Sequencing

## Abstract

Winemakers have long used copper as a fungicide on grapevine. However, the potential of copper to accumulate on soil and affect the biota poses a challenge to achieving sustainable agriculture. One recently developed option is the use of biocontrol agents to replace or complement traditional methods. In the present study, a field experiment was conducted in South Africa in which the leaves in two blocks of a vineyard were periodically treated with either copper sulphate or sprayed with *Lactobacillus plantarum* MW-1 as a biocontrol agent. This study evaluated the impact of the two treatments on the bacterial and fungal communities as they changed during the growing season. To do this, NGS was combined with quantitative strain-specific and community qPCRs. The results revealed the progression of the microbial communities throughout the season and how the different treatments affected the microbiota. Bacteria appeared to be relatively stable at the different time points, with the only taxa that systematically changed between treatments being *Lactobacillaceae*, which included reads from the biocontrol agent. Cells of *Lactobacillus plantarum* MW-1 were only detected on treated leaves using strain-specific qPCR, with its amount spanning from 10^3^to 10^5^cells/leaves. Conversely the fungal community was largely shaped by a succession of different dominant taxa over the months. Between treatments, only a few fungal taxa appeared to change significantly and the number of ITS copies was also comparable. In this regards, the two treatments seemed to affect the microbial community similarly, revealing the potential of this biocontrol strain as a promising alternative among sustainable fungicide treatments, although further investigation is required.

## 1. Introduction

Copper (Cu) compounds have traditionally been the means to combat phytopathogenic microbes, especially fungi, in crop plants (Banik and Pérez-de-luque 2017). In organic agriculture including organic vineyards, the use of Cu-based pesticides is currently the only chemical treatment allowed, although it is limited to a maximum of 6 kg Cu ha^-1^ per year in the EU (Commission Regulation, 2002). Such a strict regulation is explained by the adverse effects of Cu on soil organisms and its long-term persistence on surface horizons (Van Zwieten et al. 2004). Another rational behind this is that Cu overuse may lead to the development of Cu resistance in pathogenic fungi (Wang et al. 2011). Lastly, Cu residues on grapes may compromise wine quality (Tromp and Klerk 1988). In contrast, biocontrol agents, such as lactic acid bacteria, are not harmful to either the environment or health. Indeed, they are classified as “generally recognized as safe” (GRAS) by the Food and Drug Administration (FDA, USA) and have received the “qualified presumption of safety” (QPS) status from the European Food Safety Agency (EFSA) (Trias et al. 2008).

The biocontrol potential of the lactic acid bacterium *Lactobacillus plantarum* has recently emerged as an important subject of study. The effectiveness of *L*. *plantarum* against several fungi has been emphasized in a large-scale screening study of over 7.000 lactic acid bacteria, which showed that most of the strains with antifungal properties belonged to *L*. *plantarum* (Crowley, Mahony, and van Sinderen 2013). Furthermore, many *in vitro* studies have demonstrated the antifungal properties of *L*. *plantarum* strains against plant parasites of the genera *Botrytis* (Sathe et al. 2007; Wang et al. 2011; Fhoula et al. 2013; de Senna and Lathrop 2017), *Fusarium* (Lavermicocca et al. 2000; Sathe et al. 2007; Smaoui et al. 2010; Wang et al. 2011; Crowley, Mahony, and van Sinderen 2013), *Aspergillus* (Lavermicocca et al. 2000; Djossou et al. 2011), *Rhizopus* (Sathe et al. 2007; Djossou et al. 2011), *Alternaria, Phytophtora* and *Glomerella* (Wang et al. 2011), *Sclerotium, Rhizoctonia* and *Sclerotinia* (Sathe et al. 2007). Additionally, chili seeds infected by *Colletotrichum gloeosporioides* germinated well after treatment with *L*. *plantarum* (El-Mabrok et al. 2012). In the field, a mix of *L*. *plantarum* and *Bacillus amyloliquefaciens* applied to durum wheat from heading to anthesis showed promising control of the fungal pathogens *F*. *graminearum* and *F*. *culmorum* (Baffoni et al. 2015).

*In vitro* trials using different *L*. *plantarum* strains have shown a promising inhibition of plant pathogenic bacteria belonging to the genera *Clavibacter* (Oloyede et al. 2017), *Xanthomonas* (Visser et al. 1986; Trias et al. 2008; Roselló Prados 2016), *Erwinia* (Visser et al. 1986) and *Pseudomonas* (Tajudeen et al. 2011; Fhoula et al. 2013; Roselló Prados 2016). In fact, the inhibitory activity of *L*. *plantarum* against *P*. *syringae* in planta has long been documented (Visser et al. 1986), while that against *Rhizobium radiobacter* has only recently been reported (Korotaeva N, Ivanytsia T, and Franco BDGM 2015). Meanwhile, field sprays of *L*. *plantarum* on Chinese cabbage alleviated the severity of soft rot by *Pectobacterium carotovorum* subsp. *carotovorum* (Tsuda et al. 2016) and contributed to a reduction of fire blight by *Erwinia amylovora* on apple and pear (Daranas et al. 2018). However, the aforementioned studies focused on the effects of *L*. *plantarum* on single organisms while its effects on the microbial community in a field remain to be investigated.

Next-generation sequencing (NGS) and quantitative PCR (qPCR) techniques facilitate rapid and cost-competitive mapping of complex microbiomes by tracking fastidious, unculturable or even unknown taxa and their roles (Schloss and Handelsman 2005; Clark et al. 2018). In particular, through accurate, real-time enumeration of taxonomic or functional gene markers, qPCR has enabled microbiologists to quantify specific species or phylotypes out of an environmental “genetic soup” (Clark et al. 2018) Nonetheless, qPCR assays rely exclusively on known genes and thus overlook taxa distinct from those already described (Smith and Osborn 2009). This limitation is circumvented by NGS technologies and the approaches of either marker gene amplification (amplicon sequencing) or total (shotgun) sequencing of environmental DNA. Owing to the PCR amplification of specific molecular gene markers, amplicon sequencing can still be quite inefficient for drawing conclusions about the genus or species level (Escobar-Zepeda, De León, and Sanchez-Flores 2015). Moreover, biases can be introduced due to horizontal gene transfer, interspecific gene copy number variation within a microbiome and underrepresentation (Escobar-Zepeda, De León, and Sanchez-Flores 2015). Nevertheless, both NGS and qPCR have substantially contributed to explorations of the planet’s microbiome, and amplicon sequencing remains a valuable tool for comparative studies of microbial communities (Warinner et al. 2014; Roggenbuck et al. 2014).

From an anthropocentric viewpoint, there has been increasing interest in using the above technologies to decipher the microbial interactions that directly affect human health and resources (*e*.*g*. on crops and livestock). In this context, the positive role of probiotic bacteria has long been acknowledged and their potential is being investigated in some detail (Soccol et al. 2010). *L*. *plantarum* is a versatile lactic acid bacterium and probiotic, and is ubiquitous on plants. The bacterium has been isolated from various environments, such as fermented food products, the human mouth and grape must (König and Fröhlich, 2009), and hence constitutes a promising case study. *L*. *plantarum* has been examined for, among other things, its adequacy as a biocontrol agent against phytopathogens and, as mentioned previously, it is generally considered a good candidate for use in agriculture.

Even though available results of the antagonism of *L*. *plantarum* against phytopathogens *in vitro* are encouraging, harsh field conditions call for the careful design of field application trials (Visser et al. 1986). For example, a lack of nutrients can be circumvented by repeated, surface spray applications to sustain high viable biocontrol numbers (Visser et al. 1986). The scarcity of field studies makes it more difficult to evaluate this prediction, as well as the effect *L*. *plantarum* may have on a field’s native microbiome. The aim of this study was to evaluate this effect by investigating the progression of the bacterial and fungal communities in two plots in the same vineyard, one treated with Cu and the other with a biocontrol strain of *L*. *plantarum*, as the season progressed. Furthermore, the persistence of the applied biocontrol organism on the vines was monitored. To the authors’ knowledge, this study is the first preventative intervention using a biocontrol *L*. *plantarum* strain in a field-experiment.

## 2. Results

This study used a combination of amplicon library NGS for both 16S and ITS, and qPCR with universal and strain-specific primers. The outcome of the DNA sequencing is presented below, followed by the results of qPCR analyses on the vine leaves. Finally, the results are presented of analyses of the must during different steps in the fermentation process.

### 2.1 DNA sequencing dataset description

The sequencing provided a dataset of 147 samples. After quality filtering, 14,028 exact sequence variants were attained from 3,011,656 high-quality reads concerning just the 16S community on leaves. For the ITS sequencing of the leaves, 6,901 features were retained, coming from 4,854,508 reads. All the samples were sequenced with sufficient coverage to unravel the complexity of the microbial community harboured on the leaves, as seen by the rarefaction curves in Figure S1. To minimise statistical issues that could arise from differences in terms of sequencing depth, the samples were normalised by randomly extracting 24,000 reads from each of the samples before any downstream analyses were performed.

### 2.2 Alpha diversity on leaves

This section focuses on two different parameters that describe the microbial community on leaves: phylogenetic diversity (PD) and evenness. The alpha diversity results are summarised in Figure 1. Looking at the bacterial community on leaves, it was noted that PD was significantly higher in the Cu-treated samples compared to the biocontrol samples, as shown in Figure 1a (p-value= 0.014), and that PD seemed largely unaffected by the sampling time during the season (p-value= 0.538), as shown in Figure 1c. Looking at the evenness in Figure 1a, it can be seen that the Cu samples had a greater evenness than the biocontrol samples (p-value =0.0003). In contrast, no statistical effect on evenness was evident when looking at the collection time for 16S (p-value= 0.38), as displayed in Figure 1c.

**Figure 1:**
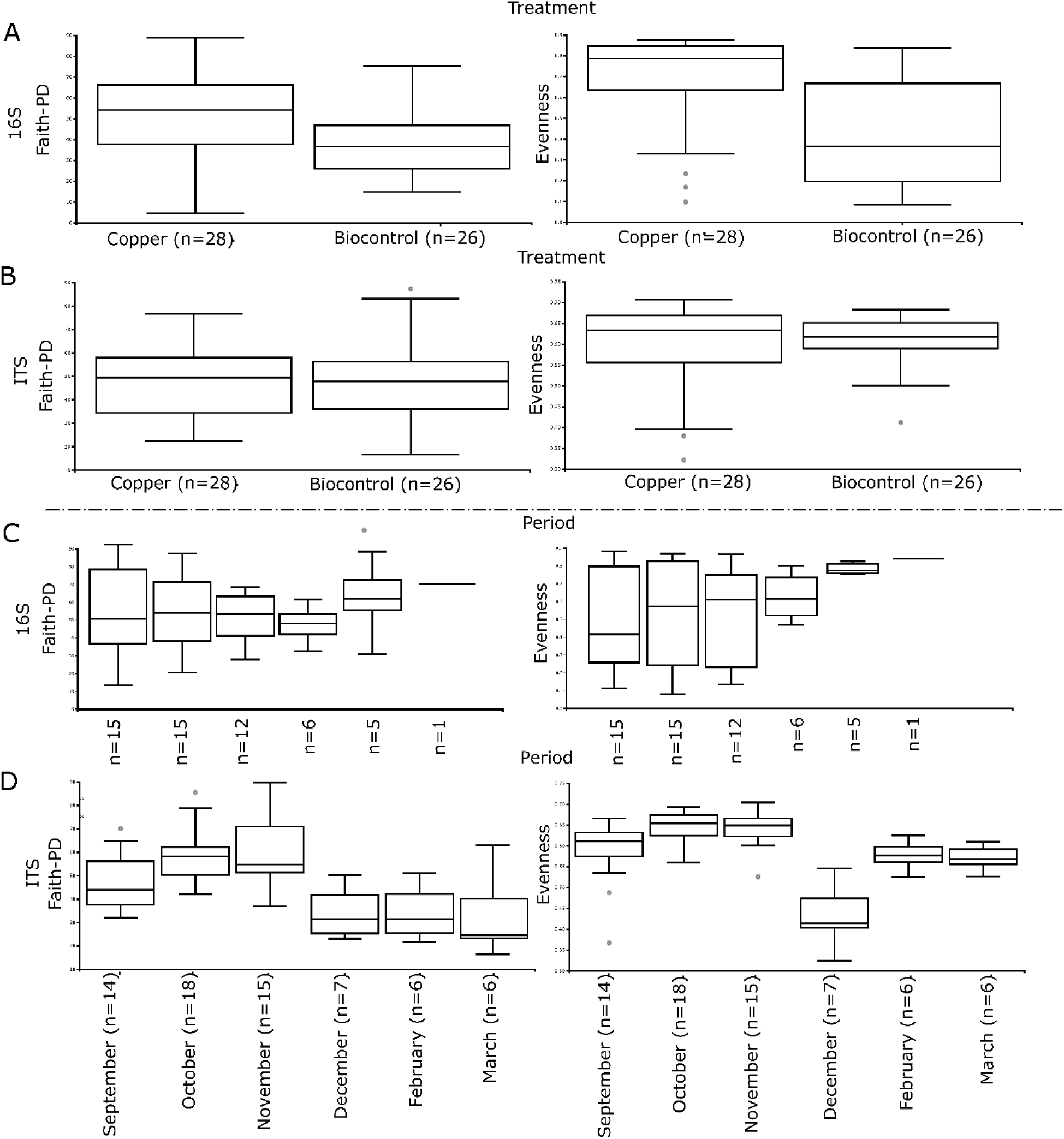
Boxplots of phylogenetic diversity (PD) (left) and evenness (right), grouped by treatments (above) and period (below) for bacteria and fungi. a) PD (left) and evenness (right) of bacteria grouped by treatment, b) PD (left) and evenness (right) of fungi grouped by treatment, c) PD (left) and evenness (right) of bacteria grouped by period, d) PD (left) and evenness (right) of fungi grouped by treatment.

In contrast, the fungal community seemed to be unaffected by the treatments in terms of PD and evenness (p > 0.05), as visualised in Figure 1b. However sampling time produced a strong effect on PD (p-value<<0.05) and evenness (p-value<<0.05) on the ITS distribution, as revealed in Figure 1d. In particular, the number of sequence variants in the community tended to increase over time from September to November, and then suddenly decreased from December to the post-harvest period in February and March.

### 2.3 Beta diversity on leaves

This section shows the differences between the samples using Bray-Curtis dissimilarity, looking at the distribution and relative abundances of the single feature between samples. The beta diversities of 16S and ITS sequence datasets, coloured by treatment or collection time, were visualised by PCoA plots and are shown in Figure 2. With regard to the bacterial community in Figure 2a, it can clearly be seen that the 16S distribution on leaves strongly clustered by treatment (p-value=0.001), while no significant pattern (p-value>0.05) was found in relation to the sampling collection time (Fig. 2b). The opposite trend was apparent for the fungal community. In fact, although ITS distribution does not appear to be affected by the treatment (p-value >0.05) as displayed in Figure 2b, it does strongly depend on the period of collection during the season, not only at month level (p-value= 0.001) as shown in Figure 2d, but sometimes even at day level (p-value= 0.001), as presented in Figure S2.

**Figure 2:**
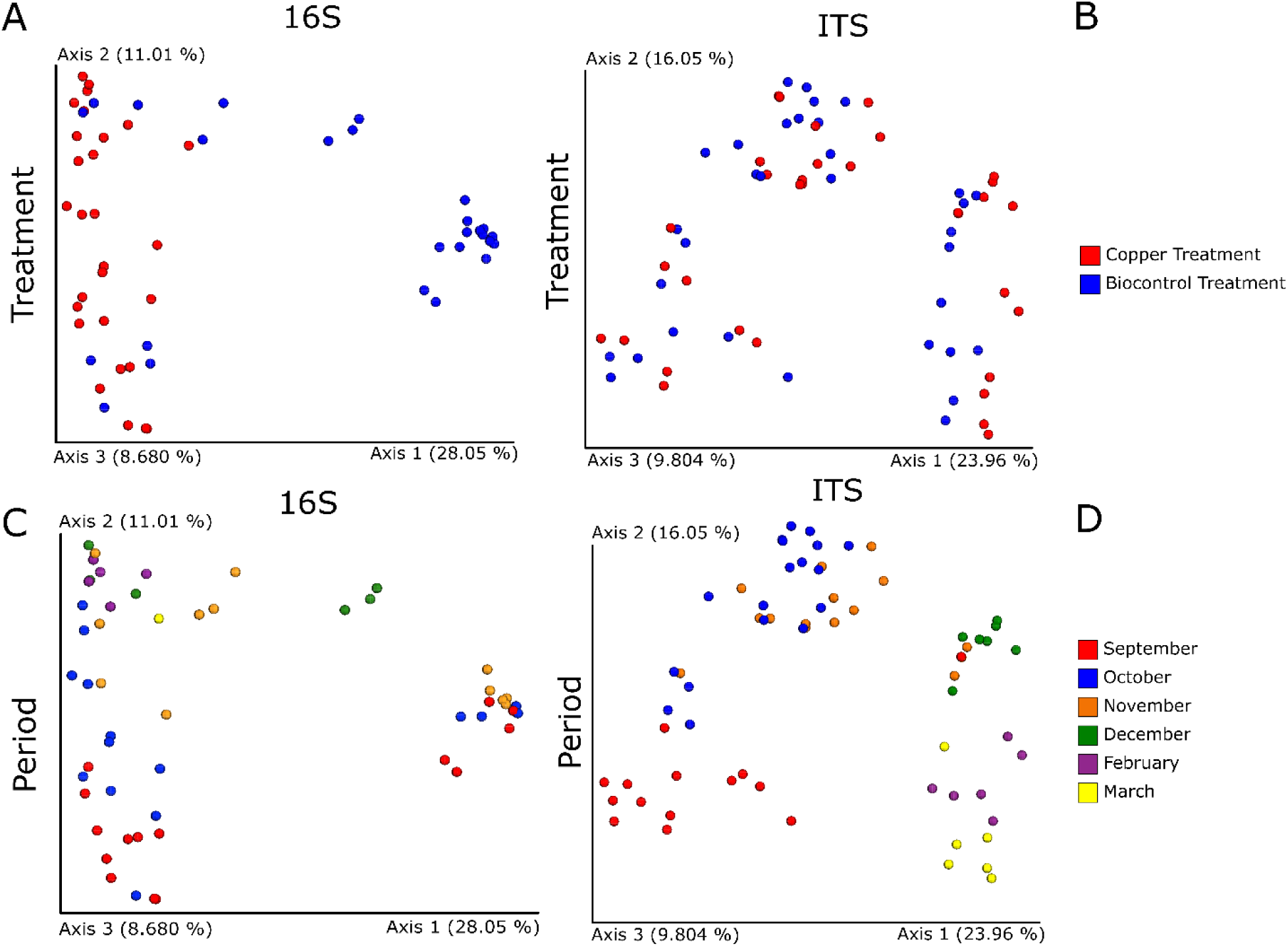
PCoA plots displaying the beta diversity within the dataset for 16S (left column) and ITS (right column) with samples coloured by treatment (above) and period of collection (below). a) Distribution of 16S data based on treatment, b) distribution of ITS dataset grouped by treatment, c) distribution of 16S dataset coloured by period, and d) distribution of ITS coloured by period. All distance matrixes are calculated based on Bray-Curtis dissimilarity.

### 2.4 Taxonomical composition on leaves

The results below show the microbial composition in terms of taxa assigned to the different features found, highlighting the microbial representation on grapevine leaves. All the information regarding bacterial and fungal community are summarised in Figure 3. Figure 3a shows the taxa bar plots of the bacterial population. In order, the dominant bacterial taxa inferred from the sequence features were *Lactobacillaceae, Bacillus, Oxalobacteraceae, Enterobacteriaceae, Planococcaceae, Pseudomonas, Enterococcus, Sphingomonas* and *Staphylococcus*. The main notable difference between the biocontrol-treated samples and the Cu-treated leaves was in the variation of *Lactobacillaceae*. This family includes the potential biocontrol agent and is an important indicator of its presence on leaves. Although members of *Lactobacillaceae* are generally present on the leaves, the difference between the biocontrol and Cu-treated samples was statistically significant (based on ANCOM). When the exact sequence variant assigned to this taxa was identified, an alignment was performed against the NCBI database using BLAST. The result of this confirmed the sequence belong to *Lactobacillus plantarum*. The presence of *Lactobacillaceae* on treated samples varied from 42 % to 68 % of the total bacterial community in the period from September to December. In the post-harvest months (February and March), the relative abundance decreased to between 0.1 % and 3.8 %, resembling the values found in the control/Cu-treated samples. This means that three months after the final spray (December) the relative abundance of *Lactobacillaceae* resembled the levels found on copper-treated leaves.

**Figure 3:**
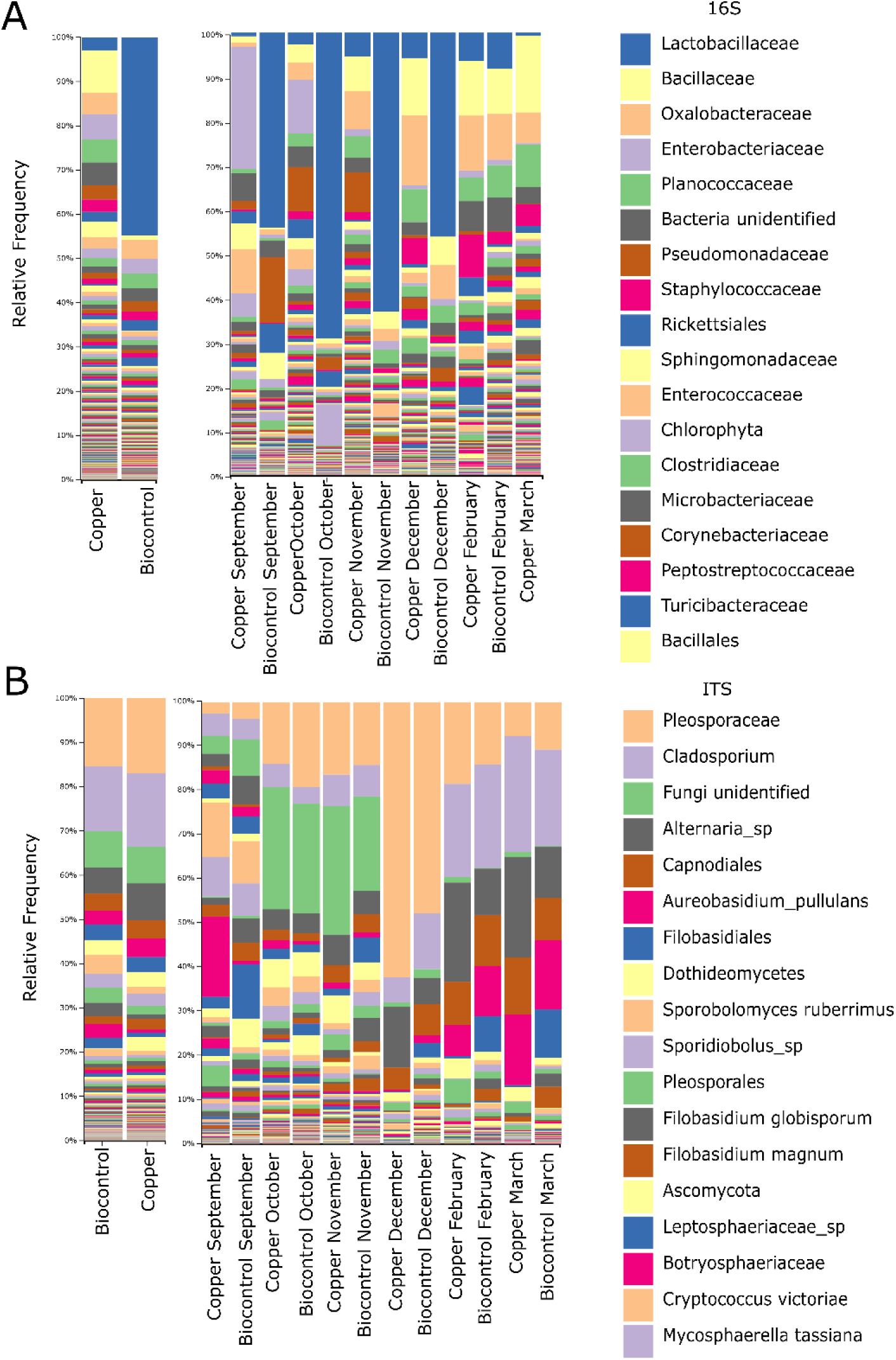
Taxonomical bar plots for the bacterial and fungal community. Figure 3a) from left to right: taxa bar plots in which all the 16S samples are grouped by treatment, a taxa bar plot in which the 16S samples are grouped by treatment during a specific period, and finally the legend representing the 18 most abundant bacteria. Figure 3b) from left to right: taxa bar plots in which the ITS samples are divided by treatment, a taxa bar plot in which the ITS samples are grouped by treatment within a specific period, and finally the legend representing the 18 most abundant fungi.

The fungal communities, displayed in Figure 3b, were relatively similar for the biocontrol and Cu-treated samples when looking at the entire growing season. The dominant taxa belonged to *Pleosporaceae, Cladosporium, Alternaria, Capnodiales, Sporobolomyces* and *Aureobasidium pullulans*. Although no differences appeared between the two treatments, another pattern was seen with respect to the collection time. In fact, during the growing season, there was a significant change in the taxonomical composition of the fungal communities. Several fungal outbreaks led to a variation in relative abundances during the period studied. For instance, *Pleosporacea* members tended to increase in relative abundance from September to December, reaching 55 % of the total community, before suddenly decreasing to 9 % in March, although they still appeared among the dominant taxa. *Sporobolomyces, Sporidiobolus* and *Botryosphaeriaceae* were more dominant in September, while *Cladosporium* and *Alternaria* were mostly represented in the post-harvest months. In light of this finding, it was of interest to examine the differences between the treatments just within the same sampling period. Interestingly *Botryosphaeriaceae* varied in relative abundance from 18 % in the Cu-treated leaves to 0.67 % to the biocontrol in September. Further analyses were performed using BLAST on the dominant sequence variant assigned to *Botryosphaeriaceae* and the read was assigned at species level to *Diplodia seriata*, a known pathogen that causes bot canker on grapevine. Conversely *Leptosphaeriaceae* detected in September ranged from 2.7 % in Cu-treated leaves to 12 % recorded in biocontrol-treated samples for the same period.

### 2.5 ANCOM and Gneiss on leaves

To test which taxa change in a statistically relevant way, ANCOM was run using treatment and period as discriminants for these samples. The result was that for 16S between different treatments, the only taxa that changed in a statistically significant way was *Lactobacillaceae*, the family to which this study’s biocontrol agent belongs, with the main sequence variant assigned to *Lactobacillus plantarum* (Table S1). Only two statistically relevant differences were found when looking at the period regarding *mitochondria* and *Ralstonia*, but none were associated with grapevine and furthermore they appeared in low abundance (Table S2).

In the fungal community there were only two taxa that changed significantly between treatments. These taxa belong to the species *Kondoa aeria* and to the order of *Filobasidiales* (Table S3). They were in very low abundances and did not seem to be related to any known disease or have an impact on the plant itself. Instead, for this period a large amount of taxa were observed that changed during the season, 42 of which are classified at least at class level (Table S4). These findings provide information that can be used to understand the evolution of the fungal community as the season progresses. These 42 taxa included potential pathogens such as *Alternaria, Cladosporium* or members of *Botryosphaeriaceae*, with the main sequence assigned to *Diplodia seriata* using BLAST. As the period shaped the fungal community, an investigation was carried out on which taxa changed between different treatments during specific periods in the season. This ANCOM test returned 50 taxa, of which 46 were classified at order level (Table S5). Among the taxa that appeared to be differentially distributed between treatments, in a short period of time (such as September alone), there were *Botryosphaeriaceae* and *Leptosphaeriaceae,* which were previously highlighted when looking at the taxonomical composition in Figure 3b.

Gneiss was then run on the bacterial community dataset to evaluate the impact of MW-1 on other bacteria. Interestingly, *Lactobacillacea* appeared to be sensitively different (p-value <0.05) from the rest of the microbial community, based on the position occupied in the tree (Fig. S3), but the fact that this taxon branches out from the rest means that it does not interact with other species in the community within the same microbial niche. This is a further evidence of that the introduction of this biocontrol agent does not alter the bacterial community that would normally be found on leaves sprayed with copper.

### 2.6 Quantifying fungal and MW-1 biomass on leaves

The results of the quantitative evaluation by qPCR of the microbial community on the samples are summarised in Figure 4. The detection of the biocontrol agent was determined by strain-specific qPCR combined with a high-resolution melting curve. Furthermore, the total relative abundance of fungi during the period and between the treatments was compared using universal primers for fungi (with the same primer set used for the generation of the sequencing reads to reduce biases). The biocontrol agent was detected in all the treated samples after the first spray (26 September) and up to 2 February, one month after harvest. The cell numbers showed a range spanning from 10^3 to 10^5 cells/leaf. The maximum amount of *Lactobacillus* plantarum MW-1 cells was detected after the spray on 15 November, while on two dates the number of cells detected was below the detection limit. This study’s time zero, 15 September, was the first time that MW-1 was not detectable, with no spraying having previously been done, while the second time was on 9 March, two months after harvest.

**Figure 4:**
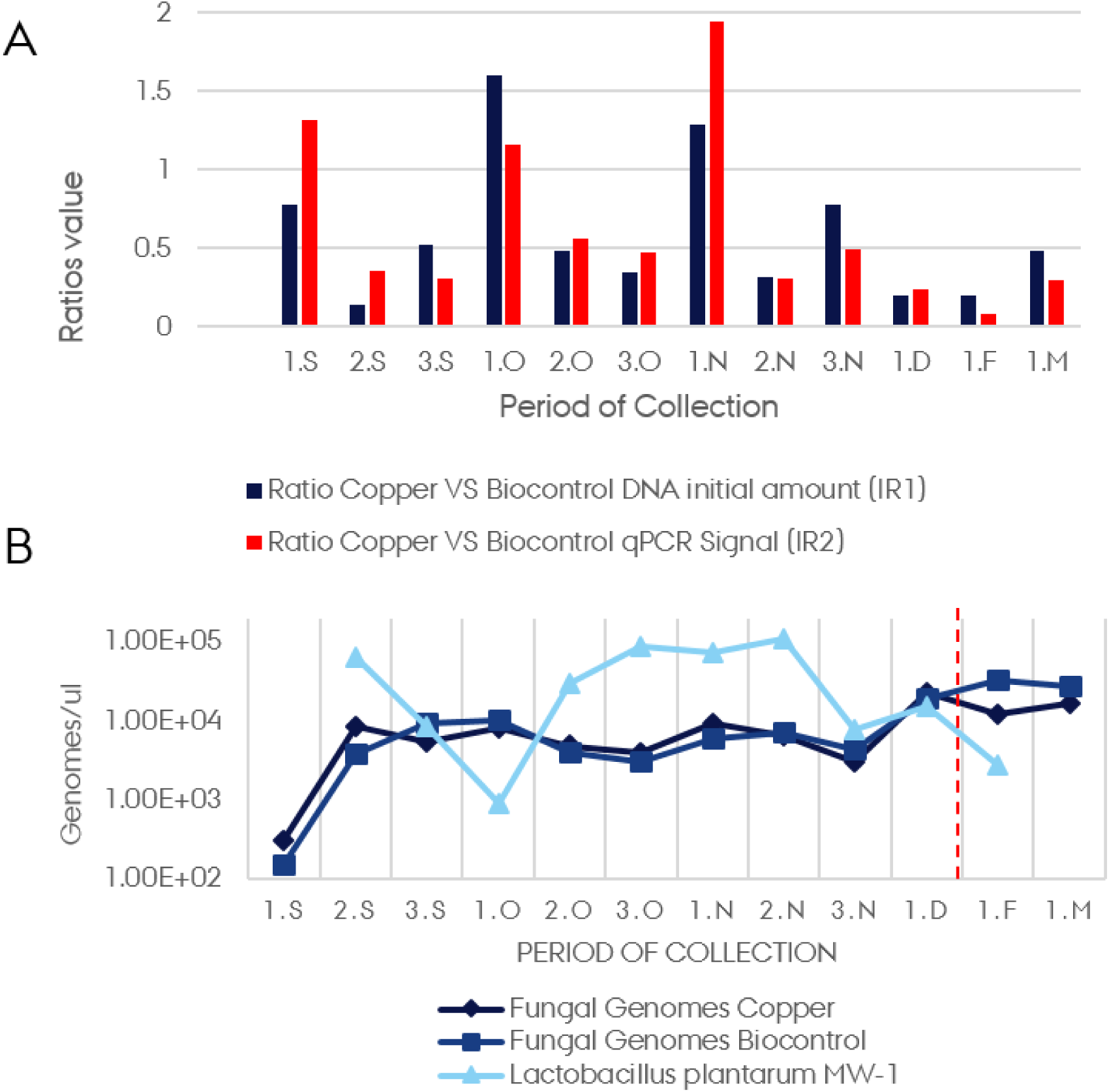
a) Positive correlation between index-ratios IR1 and IR2; letters identify the collection date as reported in Table 1. b) Normalised genomes count for MW-1 (light blue triangles), estimated fungal genomes on community treated with biocontrol agent (dark blue squares) and fungal community genomes count under Cu treatment (black diamonds); curves are expressed as log10 of the genome count. The red dashed line indicates the harvest.

The total fungal abundance could be estimated at cell level number without introducing significant biases due to an uneven ploidy variation between the different taxa detected. However, since the community composition between the treatments appeared to be very similar, it was assumed to have a negligible difference in terms of genome distribution. For this reason, a relative quantification was performed between the two fungal communities grouped by treatments. With this in mind, the results obtained showed that the fungal communities between the two treatments followed a similar trend, with no profound variations between the different treatments. It was decided to estimate the number of fungal genomes, assuming a fungal-genome average size of 27.5 Mb. Accordingly, the number of genomes ranging from 10^2 to 10^4 genomes/leaves during this period was calculated. After normalising the raw-qPCR signals using the gDNA amount, an ANOVA was run which gave a p-value of 0.56. This confirmed that the variation between the two fungal communities coming from the two treatments was not significant. Finally a correlation analysis was performed between the number of cells of MW-1 detected and the corresponding fungal genome amount for each sample. In this case, the analyses returned a coefficient of −0.39, which could be interpreted as a moderate negative correlation between fungi and MW-1.

**Table 1:**
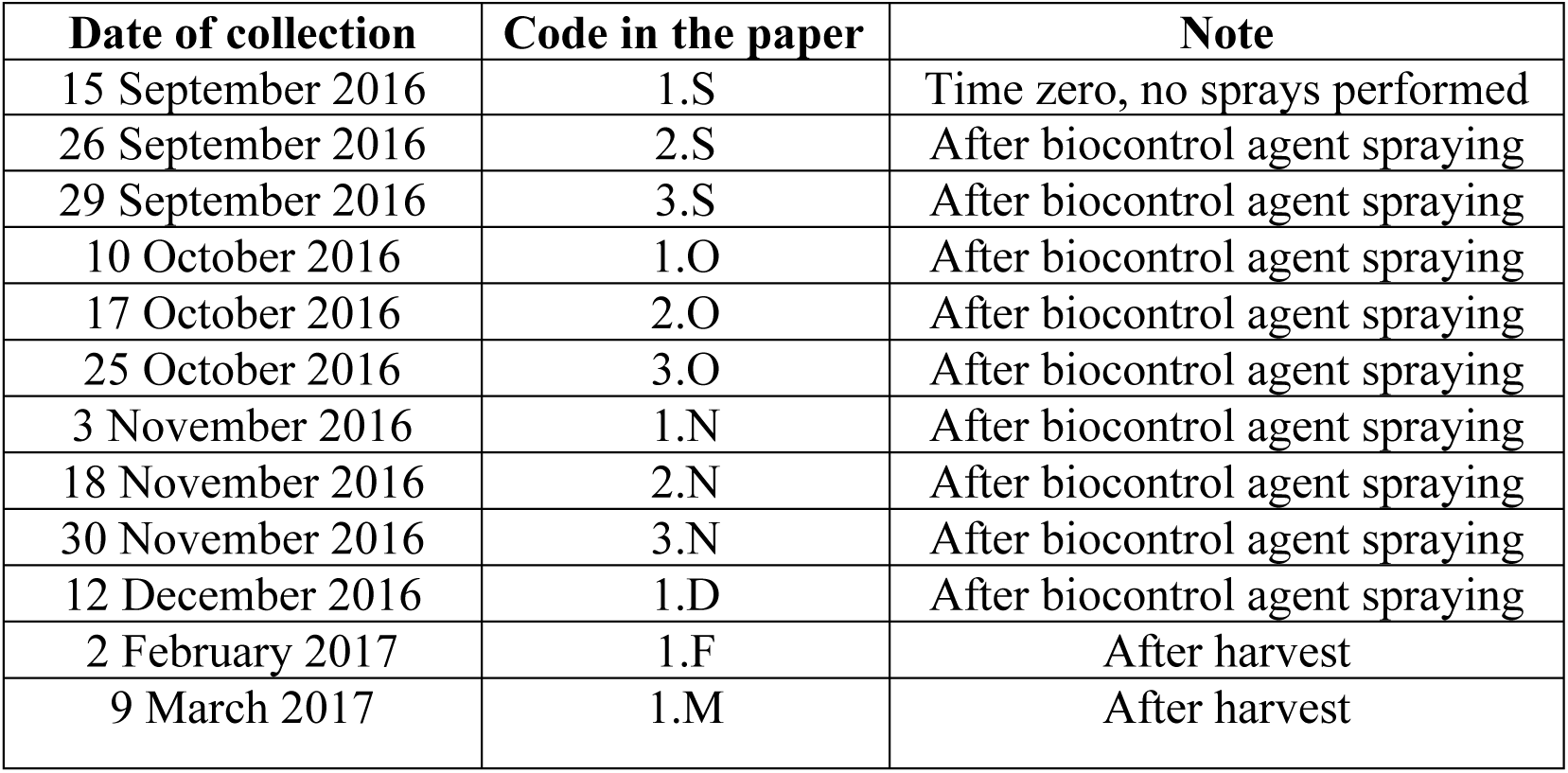
Sampling dates in this study; second column report the code used during data-processing and analyses; third column identify the state of the vineyard at the time of the samples collection

To further demonstrate the link between initial gDNA amount and qPCR signal, an additional correlation analysis was performed. First two index ratios (IR) were calculated: IR1 between input gDNA in Cu *versus* biocontrol-treated leaves, and IR2 by dividing qPCR genome-copies detected on Cu *versus* biocontrol-treated leaves. Finally, the correlation between IR1 and IR2 during the period was determined. The correlation coefficient was 0.81, confirming that the two IRs were highly positively correlated, as shown in Figure 4a. A t-test on the two IR series (the assumption of equal variance was tested with a F-test that resulted in a p-value of 0.165) confirmed that the difference was not significant (p-value= 0.699). This means that the tendency of the two IRs did not differ significantly between the treatments. In conclusion, this indicated that not only did the two fungal communities not differ significantly in composition between treatments, but they were also quantitatively comparable.

## 3. Discussion

Understanding the structure and dynamics of microbial communities in the phyllosphere is important because of their effects on plant protection and plant growth. Since the structure and dynamics of the microbiome are affected by environmental factors such as geography (Bokulich et al. 2014), grape cultivar (Singh et al. 2018) and climate (Perazzolli et al. 2014), this study focused on different treatments applied in two blocks of the same vineyard. Both treatments, Cu and MW-1, were applied to control fungal diseases. The treatments were applied throughout the growing season and were followed by the periodic collection of samples. The number of sprays was unequally distributed and was based on the visible symptom evaluation on the plants within the two blocks. This is important to note since the impacts of any chemical and biological treatments depend on the dosage and frequency of applications, as well as other biotic and abiotic factors (Perazzolli et al. 2014).

This study revealed the development of the grapevine leaf microbiome over the growing season and displayed the impact of different types of vineyard management using NGS technology combined with qPCR. The combined use of these approaches allowed quantitative and qualitative information to be recovered from the compositional dataset (Perazzolli et al. 2014).

In the present study, the biocontrol-treated phyllosphere-bacterial community had a reduced phylogenetic diversity and evenness compared with Cu-treated leaves, seemingly due to the presence of highly dominant taxa (*i*.*e*. the biocontrol agent, which appears as *Lactobacillaceae*, being dominant in almost all the treated samples). These highly abundant taxa probably compact the remaining populations, reducing evenness and phylogenetic diversity, and excluding some of the rare species that are likely to be missed at a fixed sequencing depth. In fact, phylogenetic diversity positively correlates with sequencing depth, as shown by Davison and Birch (2008). The beta diversity of the bacteria displayed significant differences since both communities, by treatment, scattered in two distinct directions along axis 1, which explained 28.5 % of the variation.

In contrast to the bacterial communities, the fungal communities were affected by the sampling period, during which PD and evenness showed similar trends for both the biocontrol and Cu-treated leaves. This is in line with Singh et al. (2018) who showed that the shift of the microbial community in samples collected at different times was detected for fungi only. The fact that phylogenetic diversity increased, together with evenness, until December suggested that the complexity of the fungal community increased due to a higher number of equally abundant taxa. In December, the fall in PD and evenness was probably due to the presence of one or a few taxa with a high relative abundance that replaced rare species. This hypothesis is based on the fact that the sequencing depth was comparable between samples. The beta diversity of the fungal communities showed that the two treatments had an insignificant effect on the fungal community when the whole period was considered. This result is consistent with the study of Perazzolli et al. (2014) where a chemical and a biocontrol agent against downy mildew were tested in different locations and climatic conditions. They showed that the fungal community was impacted mostly by geography and was resilient to the treatments tested. Instead, it is clear from the present study that the fungal community was strongly affected by the sampling time, suggesting an evolution of the harboured microbiota due to seasonal changes such as temperature, solar exposition, rainfall and other abiotic fractions. The fact that microbial seasonal shift only impacts the fungal community and not the bacterial community is consistent with the work of Singh et al. (2018) in which two sets of samples from the same vineyards were analysed and the impact of different sampling times on the microbial community was measured.

The differences in beta diversity reflected the change in taxonomical composition in both bacterial and fungal communities. In particular, the bacterial communities differed significantly only for the relative abundance of *Lactobacillaceae,* as shown in Table S1, obtained by applying ANCOM. This taxon appeared in the biocontrol-treated leaves in relative abundance up to 40 % and contained reads of the biocontrol agent MW-1. After blasting the exact sequence variant against NCBI, the analyses could be refined to species level, identifying *Lactobacillus plantarum* as the most likely assignment. The present biocontrol strain, which here appeared as the dominant sequence variant, was not the only representative of *Lactobacillaceae*. The presence of other sequence variants assigned to *Lactobacillaceae* was also retrieved on copper-treated samples. This is consistent with the existing literature in which a community of LAB is found living on the phyllosphere naturally (Daranas et al. 2018). Furthermore, the strain-specific qPCR applied to recognise and quantify MW-1 produced a precise signal peak only on the biocontrol-treated samples. This allowed the conclusion to be drawn that the different amount detected in relative abundance for the *Lactobacillaceae* family was due to the presence of reads from the biocontrol agent. Finally the bacterial community, which appeared to be relatively stable throughout the season, reflected the composition already reported by Singh et al. (2018) The dominance of Proteobacteria and Firmicutes highlighted in their paper was consistent with the distribution shown in the present study, where the dominant taxa belonged to the same phyla. Furthermore, Singh et al. (2018) analysed two different sets of samples, one collected in spring and one during the harvest period, which displayed similar bacterial community composition with a few differences in relative abundance, *e*.*g*. the taxa *Cyanobacteria*.

The analysis of the microbial community throughout the season offered interesting insight into the possible correlations between specific fungal outbreaks and seasonal variations. When looking at the whole period, there was a clear succession of dominant fungal taxa with *Botryosphaeriaceae* and *Sporobolomyces* in September, *Pleosporacea* in December and *Cladosporium, Alternaria* and *Aureobasidium pullulans* in February and March. All of these taxa are listed as being commonly recovered on grapevine plants in the review of Dissanayake et al. (2018). However, focusing on the differences between treatments that appeared only at a specific period of the season, other taxa could be found that were sensitive to the treatments. These taxa were generally hidden when looking at the complete timeline, but appeared clear when the focus was on a specific period. For instance, in September, just after spraying started, there were differences in the dominant taxa between treatments. The biocontrol leaves showed a reduced relative abundance of *Botryosphaeriaceae* compared to the Cu-treated samples, with the dominant sequence variant assigned to *Diplodia seriata*, a known grapevine pathogen that can cause bot canker (Úrbez-Torres et al. 2008). Conversely, within the family *Botryosphaeriaceae,* there is also the genus *Botryosphaeria*, whose anamorph stage has been confused with *Diplodia* (Phillips, Crous and Alves 2007). This genus appears not to be strongly affected by Cu (Mondello et al. 2017), but from the results of the present study it might be impacted by MW-1. Conversely, this biocontrol agent seems not to affect the taxon *Leptosphaeriaceae,* whose group contains pathogenic taxa such as *Leptosphaeria* that can cause black-leg disease in brassica (Hammond, Lewis, and Musa 1985). However, to the authors’ knowledge, *Leptosphaeriaceae* does not include pathogenic strains on grapevine. Furthermore, *Leptosphaeriaceae* are sensitive to chemical pesticides (Eckert et al. 2010), and this could explain the reduction when Cu was applied. However targeted quantitative methods are required to confirm this.

Finally, the outcome of the NGS approach was combined with a targeted strain-specific qPCR to detect and quantify the biocontrol agent. A qPCR was applied on the fungal communities from the two treatments. This combined approach has previously been reported (Perazzolli et al. 2014) as an efficient way of drawing conclusions about microbial ecology when biocontrol agents are involved. The number of cells detected for MW-1 was consistent with the existing literature in which biocontrol agents on the phyllosphere are studied and quantified, and could eventually be used as an indicator of the effectiveness of the treatment (Perazzolli et al. 2014). The detection limit in this study was 10^2^ cells/gram of leaves, a frequent limit of detection when qPCR is used (Hierro et al. 2006). The number of cells detected during the season varied between 10^5^and 10^3^. A sudden tenfold decrease was expected 1-6 days after the spray, as previously reported in other field studies such as Daranas et al. (2018) using *L*. *plantarum* PM411 on apple, kiwi, strawberry and pear plants. When investigating the amount of fungi, the number of cells was not estimated due to the biases that could have been introduced by different ploidy between species. However, an average amount of fungal genomes was estimated based on the information reported in Li et al. (2018), with the average size for ascomycetes and basidiomycetes estimated at 13 and 42 Mb respectively. Accordingly, since members of both phyla were retrieved, 27.5 Mb was used as the average size for genomes. However, the results could be slightly different if these parameters were changed, consistent with other studies on fungal genomes such as Mohanta and Bae (2015), which reported different sizes. All the analyses established that the two fungal communities were quantitatively similar. Finally, when comparing the two normalised qPCR signals, an ANOVA test confirmed that there were no significant differences between treatments.

Based on these results, leaves treated with *L*. *plantarum* MW-1 showed minor differences at specific periods of the year in terms of taxonomical composition, and there were no significant differences in terms of fungal load when compared with Cu-treated leaves. To conclude, although further evidences are required, this study provides affirmation that *Lactobacillus plantarum MW-1* offers a promising biocontrol alternative or supplement to the use of Cu as a fungicide treatment on grapevine leaves. Finally, it paves the way for more environmentally-friendly treatment practices, not only in vineyards but also in other crops that are sensitive to fungal diseases and require chemical treatments.

## 4. Material and Methods

### 4.1 Materials

Leaves were collected periodically from September 2016 to March 2017 from a vineyard in a winery in the Western Cape (South Africa). Two plots in the same field, separated by several lines of grapevine, were managed in two different ways: one block was sprayed with Cu in a traditional vineyard management system, while the other was sprayed with a *Lactobacillus plantarum* MW-1 as a potential biocontrol agent. The *Lactobacillus plantarum* MW-1 strain has previously been isolated from grapes in the same wine region (data not shown). During the studied period, 12 different samplings were carried out in which leaves were collected in five spots for each of the treatment areas within the vineyards. For each spot and each treatment there were three independent biological replicates. Each sample consisted of five leaves that were placed in a 50 ml Falcon tube. The tubes were frozen at −20 °C and shipped to the laboratory in Denmark. Sample collection followed the spray calendar for the biocontrol agent. After the grape harvest in January, there was a follow-up with two extra samplings in February and March. All the plants from which leaves had been collected were marked and did not change throughout the experimental plan. A complete calendar of the sampling is given in Table 1.

### 4.2 Sample preparation and DNA extraction

The tubes containing the leaves were thawed at room temperature and 20 ml of a washing solution was subsequently added (Singh et al. 2018) and placed on a rocket inverter, applying one hour of gently shaking rotation. After this, the leaves were discarded using sterile pincettes. The washing solution was centrifuged for 15 minutes at 6000 rpm to create a pellet. Following removal of the washing solution, the pellet was resuspended in 978 ul of phosphate buffer solution and DNA was extracted according to the manufacturer’s protocol using FAST DNA Spin Kit for Soil (MP-Biochemical, CA). After the extraction, the DNA quality and concentration were measured with Nanodrop (Thermo Fisher Scientific) and Qubit^®^2.0 fluorometer (Thermo Scientific™).

### 4.3 Library preparation for sequencing

Amplicon library preparation for 16S bacterial gene was performed as described by Gobbi et al. (2018), with minor modifications hereby reported. To reduce the amount of plastidial DNA from the grapevine leaves, specific mPNA and pPNA were used with the same sequences as those recommended by Lundberg et al. (2013). Each reaction contained 12 µL of AccuPrime™ SuperMix II (Thermo Scientific™), 0.5 µL of forward and reverse primer from a 10 µM stock, 0.625 ul of pPNA and mPNA to a final concentration of 0.25 mg/mL, 1.5 µL of sterile water, and 5 µL of template. The reaction mixture was pre-incubated at 95 °C for 2 min, followed by 33 cycles of 95 °C for 15 sec, 75 °C for 10 sec, 55 °C for 15 sec and 68 °C for 40 sec. A further extension was performed at 68 °C for 10 min.

The fungal community was sequenced using the same double-step PCR approach for library preparation, but the primers in use were ITS1 (TCGTCGGCAGCGTCAGATGTGTATAAGAGACAG- GAACCWGCGGARGGATCA) and ITS2 (GTCTCGTGGGCTCGGAGATGTGTATAAGAGACAG- GCTGCGTTCTTCATCGATGC) with adapters for Illumina MiSeq Sequencing. Each reaction for the first PCR on ITS contained 12 µL of AccuPrime™ SuperMix II (Thermo Scientific™), 0.5 µL of forward and reverse primer from a 10 µM stock, 0.5 µL of bovine serum albumin (BSA) to a final concentration of 0.025 mg/mL, 1.5 µL of sterile water, and 5 µL of template. PCR cycles were the same as above, but without the step at 75 °C for 10 sec. The number of cycles was set to 40. The second PCR of fragment purification using beads and Qubit quantification were common for 16S and ITS and followed the protocol reported in Gobbi et al. (2018). The final pooling was proportional to the length of the fragment in order to sequence an equimolar amount of gene fragments for all the samples. Thus by sequencing all the samples within the same sequencing run, biases due to run variations were minimised. Sequencing was performed using an in-house Illumina MiSeq instrument and 2×250 paired-end reads with V2 Chemistry.

### 4.4 Strain-specific primer for the biocontrol agent

In order to find a region that was unique for the biocontrol agent, part of the genome was uploaded and scanned using PHASTER (PHAge Search Tool Enhanced Research) (Arndt et al. 2016) to detect phage and prophage genomes within the bacterial genome. Following this, the uniqueness was tested of an amplicon generated from one primer that targets the genome and another targeting the phage sequence against the NCBI database and a private genome database repository of the strain provider. Thus a fragment of 300 bp was obtained that did not show any hit in NCBI and only one full hit in the private genome repository. To amplify this fragment, the primers MWP3F (CATCCCAACCGCTAACAA) and MWP3R (CGCAGAAAAGGTAGCAAA) were used. These primers were tested further against other strains of *L*. *plantarum* from the authors’ collection, showing amplification only toward the biocontrol agent. To create a standard curve, the DNA was extracted from a pure culture of the biocontrol agent and serial dilutions were performed from 10^-2^to 10^-7^. Knowing the length of the fragment of 300 bp, the qPCR signal and the fact that the fragment appears only once in a genome, the number of cells of the biocontrol agent *L*. *plantarum* MW-1 was estimated for each reaction. This information was used to estimate the number of cells in leaf extract samples.

### 4.5 qPCR

All PCR reactions were prepared using UV sterilised equipment and negative controls were run alongside the samples. The qPCR with primers specific for the biocontrol agent and the fungal community was carried out on a CFX Connect™ Real-Time PCR Detection System (Bio-Rad).

The primers used for the bacterial biocontrol agent were MWP3F and MWP3R, while for the total fungal community the same primers ITS1 and ITS2 were used as during library preparation. To reduce biases the adapters for Illumina MiSeq Sequencing were also maintained in qPCR. Single qPCR reactions contained 4 µL of 5x HOT FIREPol® EvaGreen® qPCR Supermix (Solis BioDyne, Tartu, Estonia), 0.4 µL of forward and reverse primers (10 µM), 2 µL of bovine serum albumin (BSA) to a final concentration of 0.1 mg/mL, 12.2 µL of PCR grade sterile water, and 1 µL of template DNA. Since DNA concentration in some of the samples was very low (around 0.5 ng/ul), normalisation by diluting was avoided before qPCR, but the qPCR signal have been normalized using the total input DNA, previously measured by Qubit. The qPCR cycling conditions included initial denaturation at 95 °C for 12 min, followed by 40 cycles of denaturation at 95 °C for 15 sec, annealing at 56 °C for 30 sec, and extension at 72 °C for 30 sec; a final extension was performed at 72 °C for 3 min. Quantification parameters showed an efficiency of E=87.7 % and R^2^of 0.999. The same qPCR cycle conditions were applied for both bacterial and fungal qPCR. All qPCR reactions were followed by a dissociation curve in which temperature was increased from 72 °C to 95 °C, rising by 0.5 °C in each cycle, and fluorescence measured after each increment. Since the initial concentration of DNA was not homogeneous and in a few samples was below 1 ng/ul, it was decided to normalise the qPCR result using the initial concentration of total DNA, as reported in Yun et al. (2006).

### 4.6. Bioinformatics

Sequencing data were analysed and visualised using QIIME 2 v. 2018.2 (Caporaso et al. 2010) and the same pipeline described in Gobbi et al. (2018). Taxonomic assignments were performed using qiime feature-classifier classify-sklearn with a pre-trained Naïve-Bayes classifier with Greengenes v_13.8 for 16S and UNITE v7.2 for ITS. The raw data of this study will be available on ENA (Study Accession Number)

### 4.7 Statistics

Statistical evaluation of these results was performed separately for qPCR and the sequencing dataset. All statistical evaluations and visualisations regarding the qPCR dataset were performed on Microsoft Office Excel 2010, while the NGS dataset was analysed through QIIME2 v. 2018.2. The qPCR signals obtained allowed the calculation of the number of cells of the biocontrol agent detected on the sprayed leaves throughout the period, while for the fungal community it was possible to perform a relative quantification between the two different treatments of Cu and the biocontrol. For both datasets the signal obtained was normalised using the total DNA as input (Yun et al., 2006).

To relate the differences in qPCR signal on the fungal community to the presence/absence of the biocontrol treatment, the qPCR signal had to be correlated with the initial DNA amount before checking the differences and their statistical relevance. To do this two different index ratios (IR) were created: IR1 dividing the raw qPCR signal from Cu samples from that from biocontrol samples, and IR2 dividing the initial amount of DNA of Cu-treated with that of biocontrol-treated leaves. Two series of IRs were then obtained to test for equal variance with a F-test. This was intended to establish whether the two IRs, deriving from two different datasets, had a comparable variance within their samples. The F-test for variances, with a p-value= 0.16, proved that the two variances were equal and then a t-test was run on two series assuming equal variances between the IRs on the two treatments, Cu and biocontrol. A correlation analysis was also performed between the initial DNA amount and the resulting qPCR signal. Finally correlation analyses and a t-test were repeated on the samples treated with the bacterial agent as well to compare the variation between the total fungal community on biocontrol-treated leaves and the number of cells detected of the biocontrol bacteria. Sequencing data after QIIME 2 pipeline processing were statistically evaluated using the Kruskal-Wallis test for alpha and beta diversity. The resulting p-value of the alpha diversity comparison was based on two parameters (phylogenetic diversity and evenness) calculated between the different series analysed. Beta-diversity analyses were evaluated using PERMANOVA with 999 permutations. Finally a statistical evaluation of differentially abundant features was performed based on analysis of composition of microbiomes (ANCOM) (Mandal et al. 2015). This test is based on the assumption that few features change in a statistical way between the samples and hence is very conservative. To individuate microbial niches, Gneiss (Morton et al. 2017) was also applied on the bacterial dataset, using treatment as the determining variable.

## Acknowledgements

The authors would like to thank the Horizon 2020 Programme of the European Commission within the Marie Schlodowska-Curie Innovative Training Network “MicroWine” (grant number 643063) for financial support.

